# NOD1 ligand FK565 promotes atherogenesis and accumulation of NOD1^high^ smooth muscle cells in atherosclerotic lesions

**DOI:** 10.1101/2025.03.14.643082

**Authors:** Xiao-Ying Zhang, Xin-Tong Jiang, Anton Gisterå, Yanchun Ding, Layth Basbous, Peder S. Olofsson, Piotr Religa, Göran K. Hansson, Zhong-Qun Yan, Maria E Johansson

**Affiliations:** Center for Molecular Medicine, Department of Medicine, Karolinska University Hospital, Karolinska Institute, Stockholm, Sweden; Department of Physiology, School of Basic Medical Sciences, Jinzhou Medical University, Jinzhou, Liaoning, China; Health Science Center, Peking University, Beijing, China; Department of Cardiology, Second Affiliated Hospital of Dalian, Medical University, China; Polish Academy of Sciences, Magdalenka, Poland; Department of Physiology, Institute of Neuroscience and Physiology, The Sahlgrenska Academy, University of Gothenburg, Sweden

**Keywords:** NOD1, Vascular Smooth Muscle, Innate immunity, Inflammation, Atherosclerosis

## Abstract

**Aims:** Nucleotide-binding oligomerization domain-containing protein (NOD)1 is an intracellular pattern recognition receptor that initiates immune responses upon ligation of molecules such as bacterial peptidoglycan containing a D-glutamyl-meso-diaminopimelic acid (iE-DAP) moiety. NOD1 ligation has been shown to promote vascular inflammation and atherosclerosis. In this study, we investigate the functional role of NOD1 in atherosclerotic plaques and characterize the vascular cells responsible for NOD1 expression and function.

**Methods and results:** NOD1 was mainly expressed in a subtype of vascular smooth muscle cells (SMC) in human atherosclerotic lesions. In *ex vivo* cultures, human endarterectomy specimens reacted to NOD1 ligand by activation of mitogen-activated protein kinase (MAPK) pathways, leading to cytokine expression. Levels of NOD1 mRNA were higher in carotid endarterectomy specimens obtained from symptomatic patients compared to asymptomatic ones. NOD1^high^ SMC were also found in arteries of atherosclerosis-prone *Ldlr*−/− mice. Challenging these mice with a NOD1 agonist resulted in transmural vascular inflammation, severe arterial damage, accelerated atherogenesis throughout the aorta, and evidence of occlusive coronary artery disease. In rats, mechanic injury to carotid arteries promoted NOD1^high^ SMC expansion and neointima formation. *In vitro*, neointima derived NOD1^high^ SMCs responded to NOD1 ligand exposure by enhanced migration, increased iNOS^+^ cells and amplified CCL5 production.

**Conclusion:** Our findings show that NOD1 promotes vascular inflammation, vascular injury responses and atherosclerosis by acting on a NOD1^high^ subtype of SMC.

## 1. Introduction

Several lines of evidence identify a pathogenetic role for innate immunity in atherosclerosis^1,2^. Innate immune responses are initiated by ligation of pattern recognition receptors such as Nucleotide-binding oligomerization domain (NOD) like receptors (NLR) and Toll-like receptors (TLR). NOD1 and its sister molecule, NOD2, act as intracellular innate immune sensors of microbial components^3^. Specifically, NOD1 recognizes γ-D-glutamyl-mesodiaminopimelic acid (iE-DAP) in peptidoglycan of Gram-negative bacteria and certain Gram-positive bacteria^4^, whereas NOD2 recognizes a muramyl dipeptide (MDP) structure in peptidoglycan of virtually all Gram-positive and Gram-negative bacteria^5^. Upon ligation, NOD1 and NOD2 activate downstream nuclear factor kappa-light-chain-enhancer of activated B cells (NF-κB) and mitogen-activated protein kinase (MAPK)s pathways via interaction with scaffolding kinase receptor-interacting protein 2 (RIP2)^6,7^.

We have previously observed a macrophage-dependent pro-atherogenic effect of NOD2 by enhancing of macrophage uptake of low-density lipoprotein (LDL) and diminishing of macrophage efferocytosis of cholesterol^8^. This, in turn, leads to expansion of the lipid-rich necrotic core of atherosclerotic lesions.

NOD1 is ubiquitously expressed in various cell types, including arterial endothelial cells and vascular smooth muscle cells (SMC)^9–11^. Endothelial cells respond to NOD1 ligation by expressing adhesion molecules that are instrumental to leukocyte adhesion, whereas SMC generate nitric oxide in response to NOD1 agonist^10^. Consistent with these *in vitro* observations, evidence from *in vivo* studies demonstrate that NOD1 agonist induces site-specific vascular inflammation^11^ and promotes atherosclerosis in atherosclerosis-prone mice^12^. Despite these advances, little is known about the contribution of individual cells to NOD1 induced immunity in vascular inflammation and atherosclerosis.

By phenotyping of NOD1 expressing cells and functional assessment of NOD1 signaling in atherosclerotic plaques from patients and experimental animals, we have identified a distinct population of SMC with a high expression of NOD1, referred to as NOD1^high^SMC, reacting to danger signals of both exogenous and endogenous origin. This response led to enhanced cell migration, cytokine and chemokine production *in vitro*, and contributed to intimal hyperplasia and occlusive arterial disease *in vivo*.

## 2. Methods

Detailed methods can be found in the online data supplement.

## 3. Results

### 3.1 Human atherosclerotic lesion contains NOD1^high^ SMC

Coronary atherosclerosis specimens obtained during autopsy were used to characterize NOD1 localization. Immunostaining analysis of specimens from five individuals were examined and revealed that NOD1 protein was preferentially co-localized with a population of α-actin positive SMC in the subcapsular region of atherosclerotic plaques and in the tunica media (Figure 1A). This NOD1^high^SMC were also found in human carotid endarterectomy plaques with a similar expression pattern (Figure 1S). Co-localization analysis revealed that 33±5% of all NOD1 positive cells were SMC. In comparison, 40±6% of all SMC express NOD1. Further analysis of samples from the biobank of Karolinska Endarterectomy (BiKE) ^13,14^ revealed a connection between NOD1 and disease manifestations. Carotid plaque NOD1 mRNA abundance was higher in patients with a preceding stroke or transient ischemic attack (Figure 1B) compared to asymptomatic individuals.

**Figure 1.**
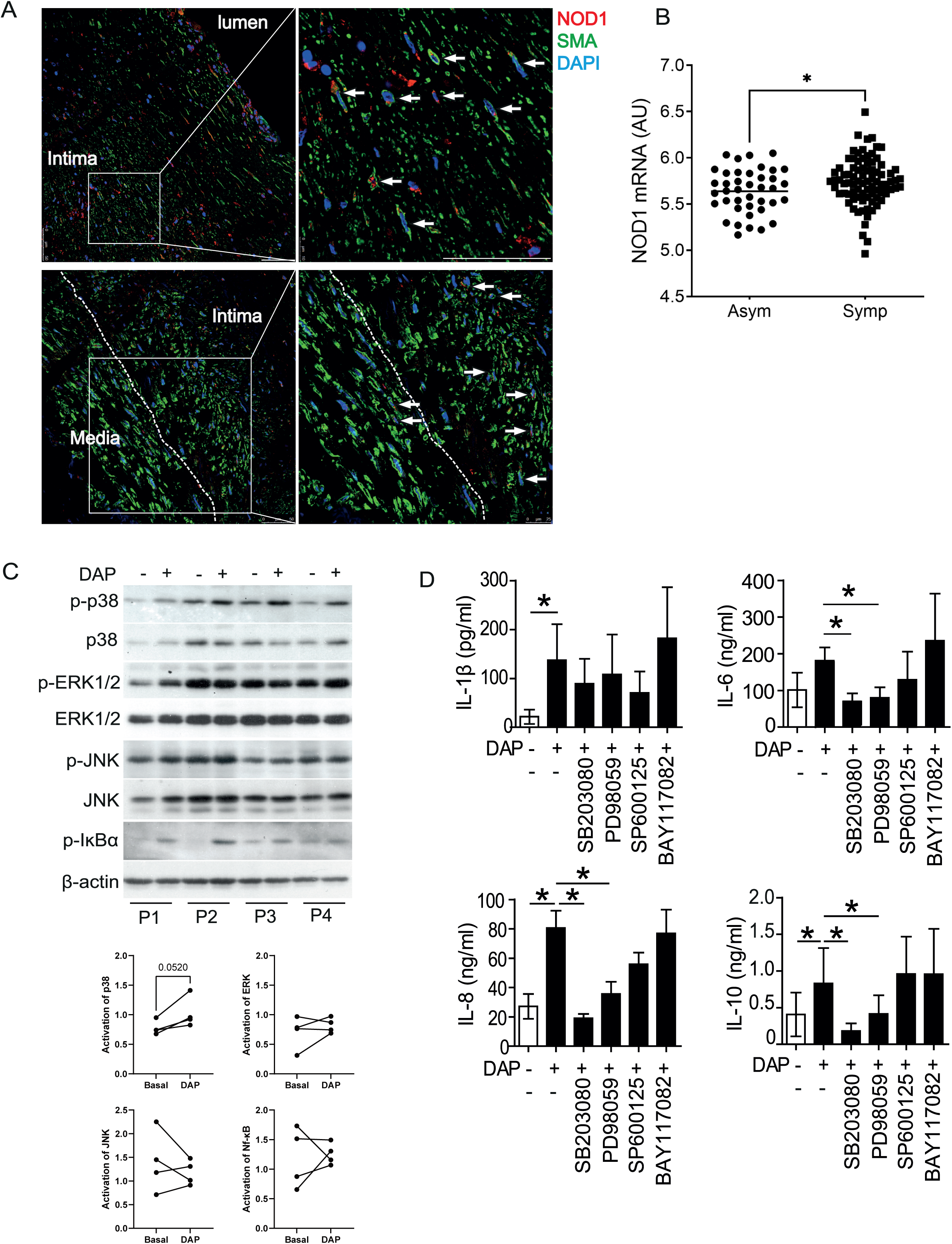
NOD1 mediates innate immune inflammation in human atherosclerosis. (A) Representative immunofluorescence micrographs of NOD1 expression in human coronary atherosclerosis lesion. Arrows indicate the cells that co-express smooth muscle α-actin (SMA, green) and NOD1 (red). Cell nuclei are labeled by 4, 6-diamidino-2-phenylindole (DAPI, blue). Scale bar, 50 µm. (B) Levels of *NOD1* mRNA in carotid plaques from symptomatic (n = 85) and asymptomatic (n = 40). Mann-Whitney, *P<0.05, data presented as scatter plots where line denotes mean values. AU, arbitrary units. (C) Top panel shows Western blot analysis of phosphorylated p38 (p-p38), p38, phosphorylated ERK (p-ERK), ERK, and phosphorylated IĸBα (p-IĸBα) in four human carotid plaques (P1-P4) treated *ex vivo* for 15 min with or without C12-iE-DAP (DAP, 1µg/ml). Lower panel shows degree of activation. β-actin was used as loading control. (D) NOD1-induced IL-1β, IL-6, IL-8 and IL-10 production by human carotid plaques pre-treated with or without p38 inhibitor (SB203080), ERK inhibitor (PD98059), JNK inhibitor (SP00125), or NF-ĸB inhibitor (BAY117082) 30 min in prior to C12-iE-DAP (DAP).

To characterize NOD1 function in atherosclerosis, we used an *ex vivo* culture system with atheromatous tissue from patients undergoing carotid endarterectomy as previously described^8^. Western blot analysis of with and without stimulation of *ex vivo* cultured atheromatous tissue with the NOD1 ligand C12-iE-DAP demonstrated activated p-38, extracellular signal-regulated kinases (ERK)1/2 and c-Jun N-terminal kinase (JNK) in all examined plaques. Sustained NF-κB activation, as detected by phosphorylated IκBα, was found in two of four plaques (Figure 1C). Moreover, *ex vivo* stimulation of atheromatous tissue with NOD1 ligand resulted in enhanced secretion of interleukin (IL)-1β, IL-8, and IL-10, while the effect on IL-6 secretion did not reach statistical significance (Figure 1D). Cytokine secretion was decreased by inhibition of ERK and p38 but appeared to be independent of JNK and NF-κB pathways.

### 3.2 NOD1 stimulation promotes experimental atherosclerosis with coronary occlusion

To examine the effect of NOD1 stimuli on atherogenesis, high fat diet fed *Ldlr*^−/−^ mice were challenged with drinking water containing the NOD1 specific agonistic ligand FK565. After ten weeks of intervention, gross examination of aorta revealed extensive atherosclerosis with outward vessel remodeling throughout the entire aorta in mice exposed to the NOD1 agonist, as compared with sham treated animals (Figure 2A). Quantitative analysis of atherosclerosis confirmed that NOD1 ligation was associated with a 3-fold enlarged lesion coverage in the aortic arch and a 2-fold increased lesion burden in the aortic root compared to the situation in mice fed high fat diet alone (Figure 2B-C). Mice subjected to the NOD1 agonist developed severe occlusive atherosclerosis in their coronary arteries (Figure 2C-D). The coronary lesions were characterized by α-actin positive SMC and CD68 positive macrophages (Figure 2D).

**Figure 2.**
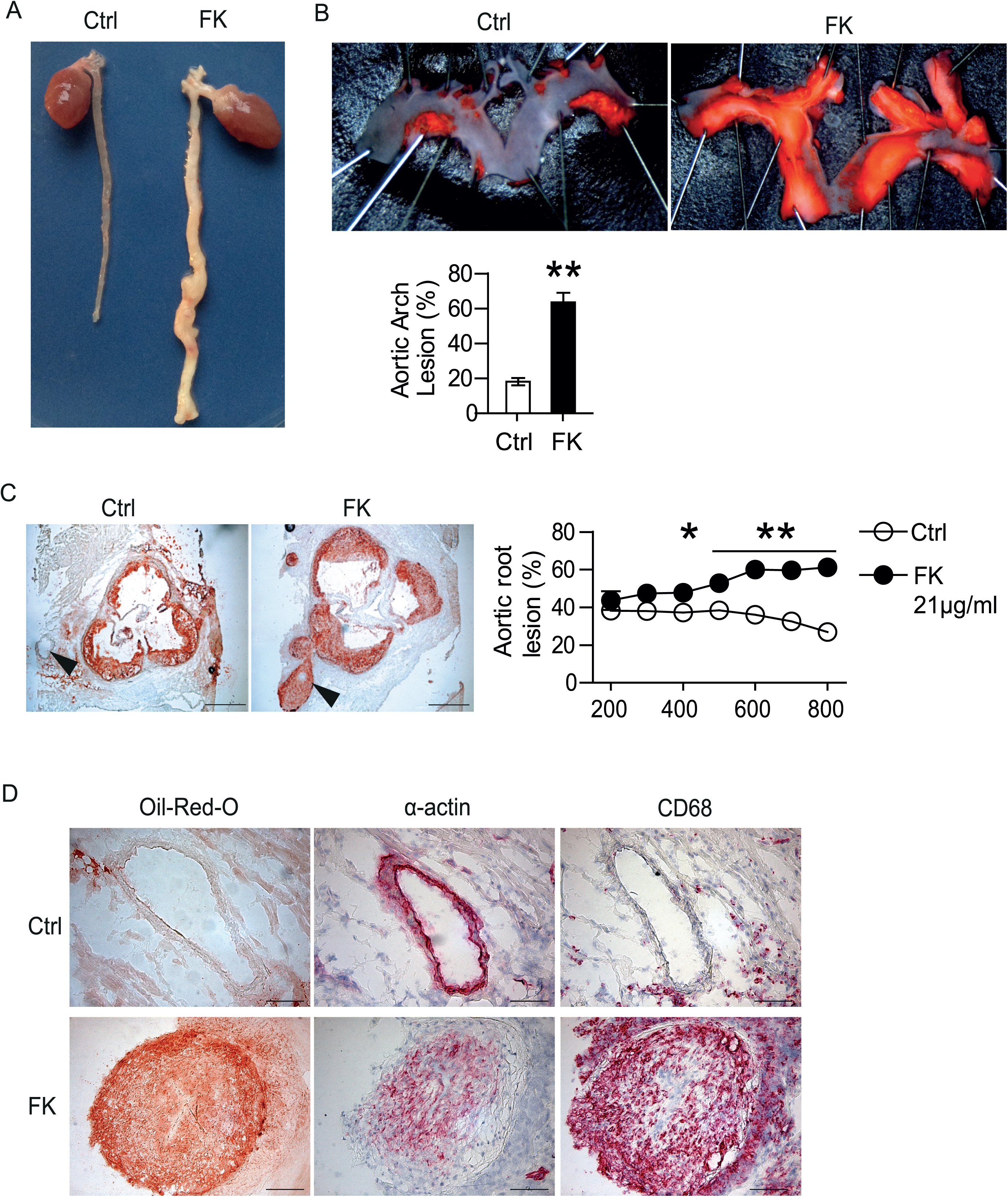
Exposure to NOD1 ligand in drinking water exacerbates development of atherosclerosis and results in coronary and brachiocephalic artery occlusion in hyperlipidemic mice. High-fat diet fed *Ldlr*^−/−^ mice were treated for 10 weeks with NOD1 ligand FK565 (FK, 21µg/ml) or without the ligand (Ctrl) in drinking water. (A) Representative gross specimens of heart and aorta in FK and Ctrl group after 10 weeks treatment. (B and C) Representative images of atherosclerotic lesion in (B) aortic arch stained with Sudan IV and in (C) aortic root stained with Oil Red O. Arrowheads indicate coronary arteries. Scale bar, 400 µm. Quantifications of atherosclerotic lesion area in aortic arch (B) and between 200 and 800 µm from the aortic root (C) are shown at right of images. Data are presented as mean ± SEM. Mann-Whitney test (B) or two-way ANOVA followed by Bonfferoni’s test (C), * p < 0.05, ** p < 0.01. (D) Representative histological analysis of coronary artery stained with Oil Red O, smooth muscle a-actin (SMA) and CD68. Scale bar, 100 µm.

In addition to the accelerated atherogenesis, 6 of 10 mice treated with FK565 died unexpectedly during the course of experiment. Despite marked differences in severity and phenotype of atherosclerosis, plasma triglycerides and cholesterol levels did not differ between the two groups of mice. However, the mean body weight of FK565-treated mice was 1.7 gram less than the control mice (p<0.05, Table S1).

### 3.3 NOD1 stimulation enhances vascular inflammation and impairs vessel wall integrity

To investigate how NOD1 ligand treatment promotes atherosclerosis, we analyzed factors known to promote vascular inflammation in atherosclerotic lesions of FK565-treated mice. FK565-treatment increased the proportion of lesion macrophages and caused upregulation of mRNA encoding nitric oxide synthase-(Nos*)2,* TNF, IL-1β, and IL-12 (Figure 3A-B, E). An increased accumulation of CD4+ T lymphocytes in the lesion was also observed, with concomitantly elevated IFN-γ and IL-17A mRNA levels (Figure 3C-E). Transcripts of key chemokines, including C-C motif ligand (CCL)2, CCL5 and C-X-C motif chemokine ligand (CXCL)10, were increased approximately 40-fold in diseased artery from the mice treated with NOD1 ligand (Figure 3F).

**Figure 3.**
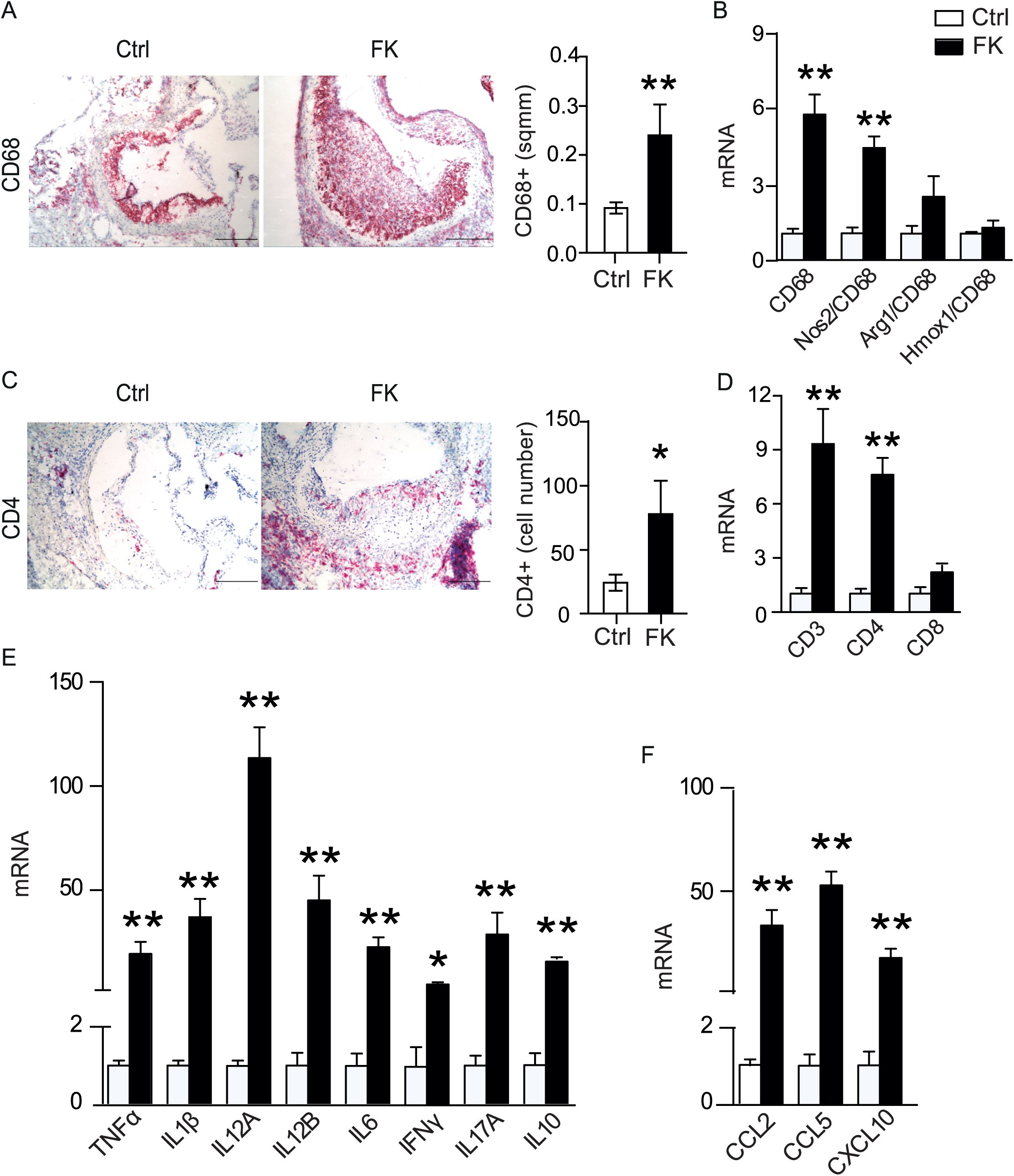
Stimulation of NOD1 enhances vascular inflammation in hyperlipidemic mice. (A and C) Representative micrographs and quantification of CD68+ macrophage and CD4+ T lymphocyte in aortic root sections from mice as in Figure 1. Scale bar, 200 µm. (B, D, E and F) qPCR analysis of (B) macrophage marker CD68 and phenotype related genes related to CD68. (D) qPCR analysis of T-cell markers, (E) cytokines and (F) chemokine transcripts in aortas. Data are presented as mean ± SEM. Mann-Whitney test was performed, * p < 0.05, ** p < 0.01.

Infiltration of CD68^+^ macrophages, Ly6G^+^ neutrophils and CD3^+^ T lymphocytes into the media was also significantly increased in FK565-treated animals (Figure 4A). Analysis of Verhoeff-stained aorta sections revealed extensive destruction of elastic lamellae in the media of NOD1 ligand treated mice. In total, 10 of 15 aortic valves in the FK565-treated mice had impaired medial integrity compared to one of 28 valves in the control group (Figure 4B). Immunohistochemical analysis showed a significant reduction of smooth muscle α-actin (α-SMA) protein in the media together with changes in medial SMC morphology in the aorta of FK565-treated mice (Figure 4C). Furthermore, relative mRNA levels of matrix metalloproteinase (MMP)9, MMP10, and MMP12 was increased in FK565-treated animals (Figure 4D).

**Figure 4.**
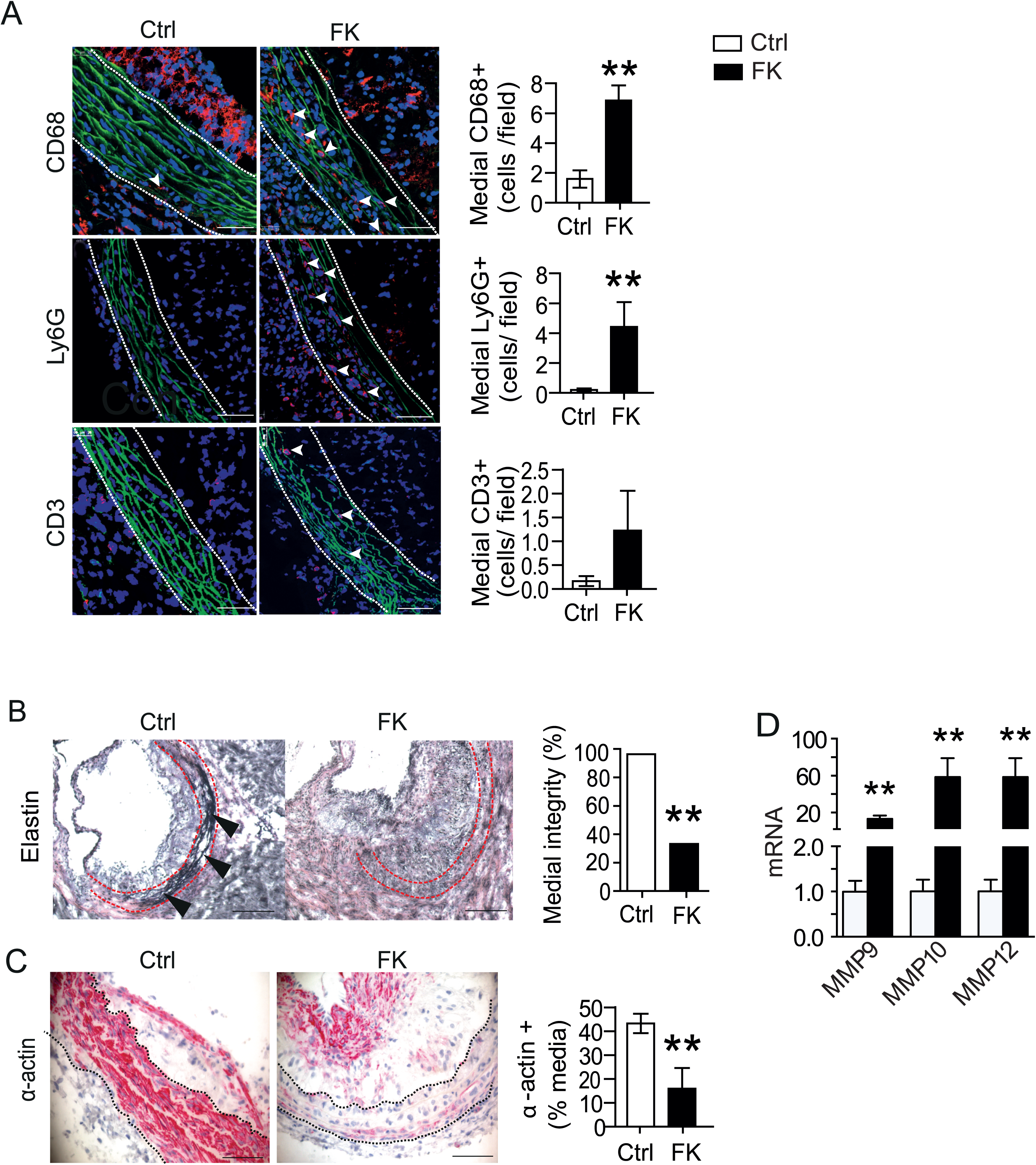
Immune cell infiltration into the arterial media layer and impaired vascular wall integrity. (A) Immunofluorescence micrographs of macrophage (CD68), neutrophil (Ly6G), and T lymphocytes (CD3) in aortic root of *Ldlr^−/−^* mice. Dotted lines mark the media layer of aorta. Scale bars, 50 µm. Bar graphs show quantification of inflammatory cells in the media. (B) Representative micrographs show Verhoeff-Van Gieson stained elastic fibers in aortic root from the mice as in Figure 1. Scale bar, 200 µm. Bar graphs show quantifications of elastic fibers in the aortic roots. Chi-square test, ** p < 0.01. (C) Representative micrographs showing immunostaining of smooth muscle α-actin (α-actin) in brachiocephalic artery sections. Scale bar, 200 µm. The graph shows the percentage of positive area for SMA in the medial layer. Mann-Whitney test, **p< 0.01. (D) qPCR analysis of mRNA encoding matrix metalloproteinase (MMP) in aorta. Mann-Whitney test, ** p < 0.01.

### 3.4 Myeloid NOD1 alone is insufficient to promote atherosclerosis

To investigate whether the presence of myeloid cell NOD1 by itself aggravates lesion size in atherosclerosis, lethally irradiated *Ldlr^−/−^* mice were reconstituted with bone marrow from *Nod1^−/−^* mice. After 13 weeks of high fat diet feeding, atherosclerotic lesions were quantified in the aortic root. There were no differences in aortic root lesion size and CD68, CD4 or TNF mRNA levels in the aorta remained unaltered (Figure 5). This implies that myeloid NOD1 is insufficient and hence vascular NOD1 required to promote atherosclerosis.

**Figure 5.**
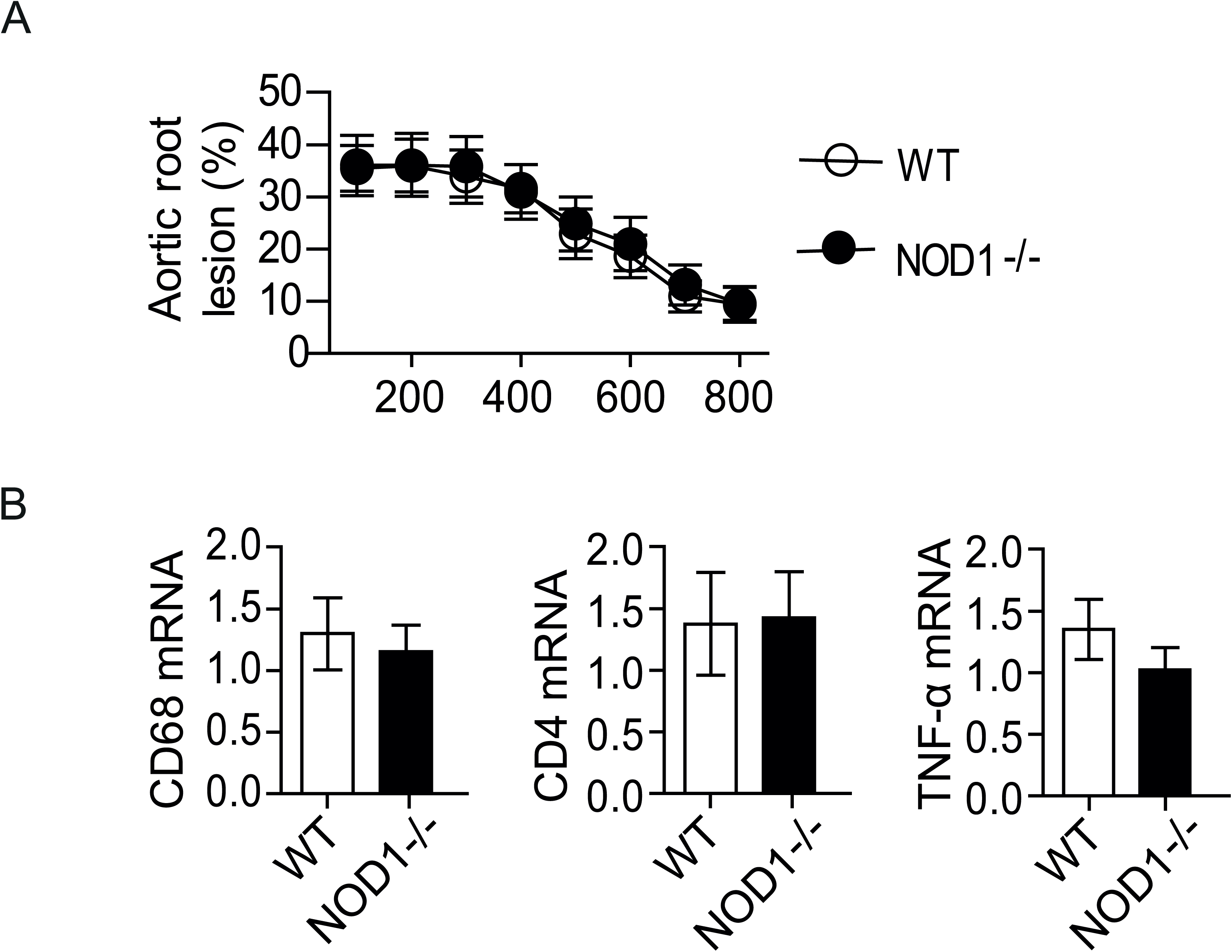
*Ldlr^−/−^* mice transplanted with *Nod1*^−/−^ bone marrow. (A) Quantifications of atherosclerotic lesion area in aortic arch stained with Oil Red O from 100 to 800 µm from the aortic root. Data are presented as mean ± SEM. (B) Gene expression of immune cell markers and the cytokine TNF in aorta quantified by qPCR. Mann-Whitney test.

### 3.5 Activation of NF-κB in NOD1^high^ SMC

Since hematopoietic NOD1 did not affect atherogenesis in *Ldlr^−/−^* mice, we proceeded to investigate cellular NOD1 expression in atherosclerotic lesions. Notably, immunostaining revealed a number of α-actin positive SMC in sections of the brachiocephalic artery samples that were distinguished from other α-actin positive SMC by expressing high levels of NOD1 (Figure 6A). These NOD1^high^ SMC were an abundant cell type in atherosclerotic lesion of mice treated with NOD1 ligand, whereas, in control mice, NOD1^high^ SMCs were predominantly present in arterial media. In fact, NOD1 expression was seen in the media in both Control and FK565-treated mice, however, the percentage of NOD1 expression in the plaques were significantly increased in FK565-treated mice (Figure 6C). Co-localisation analysis showed that NOD1 was expressed by 38±7% of all α-actin positive SMC. Furthermore, α-actin positive SMC was expressed by 46±6% of all NOD1 positive cells. Immunostaining for phosphorylated NF-κB p65, part of the canonical downstream signaling pathway of the NOD1 receptor, further showed that NF-κB activation was not present in any types of SMC in the brachiocephalic artery of *Ldlr^−/−^* control mice. However, in mice treated with NOD1 ligand, ample NF-κB activation was observed in atherosclerotic lesions and the staining was associated with regions rich in NOD1^high^ SMC (Figure 6B). These observations indicate that FK565-treatment promotes accumulation of NOD1^high^ SMC in atherosclerotic lesions.

**Figure 6.**
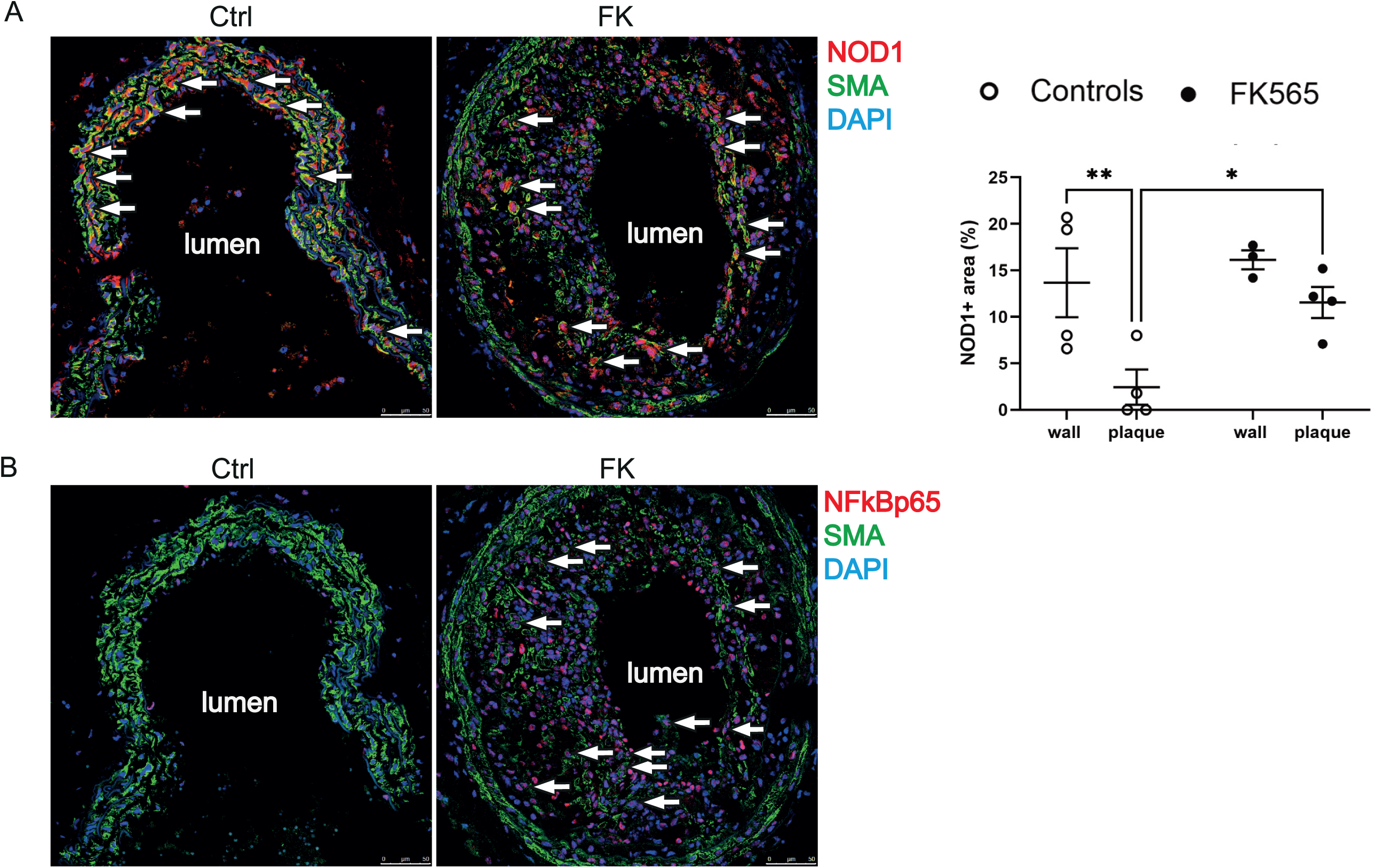
Involvement of NOD1^high^ SMC in atherosclerosis in *Ldlr^−/−^* mice. Immunofluorescence micrograph of (A) NOD1 and (B) NF-κB p65 in the brachiocephalic artery of high-fat diet fed *Ldlr*^−/−^ mice treated with NOD1 ligand FK565 (FK, 21µg/ml) or without the ligand (Ctrl) in drinking water. Arrowheads denote NOD1 or NF-κB in smooth muscle α-actin (SMA, red) positive SMC. Scale bar, 50 µm.

### 3.6 Unique response of NOD1^high^ SMC to vascular injury

Neointimal development in response to vascular injury resembles many aspects of accelerated atherogenesis and restenosis. To corroborate NOD1 function in vascular injury, alteration in NOD1 mRNA expression during the course of neointima development was assessed in a rat model of vascular injury. A time dependent increase of NOD1 mRNA was observed in carotid artery after injury. A twofold increase was observed at day 3 and a fourfold increase was observed at day 14 relative to the contralateral sham-injured carotid artery (Figure 7A). Immunostaining for NOD1 further associated the increase in NOD1 mRNA with occurrence of NOD1^high^ SMC population in the neointima (Figure 7B). Analysis of isolated SMC populations confirmed that constitutive NOD1 mRNA was threefold higher in neointima-derived SMC versus medial SMC (Figure 7C). In culture, NOD1 mRNA levels remained stable in neointima derived SMC between three and seven passages (data not shown). Thus, imprinted NOD1 expression seems to be a hallmark of neointimal SMC.

**Figure 7.**
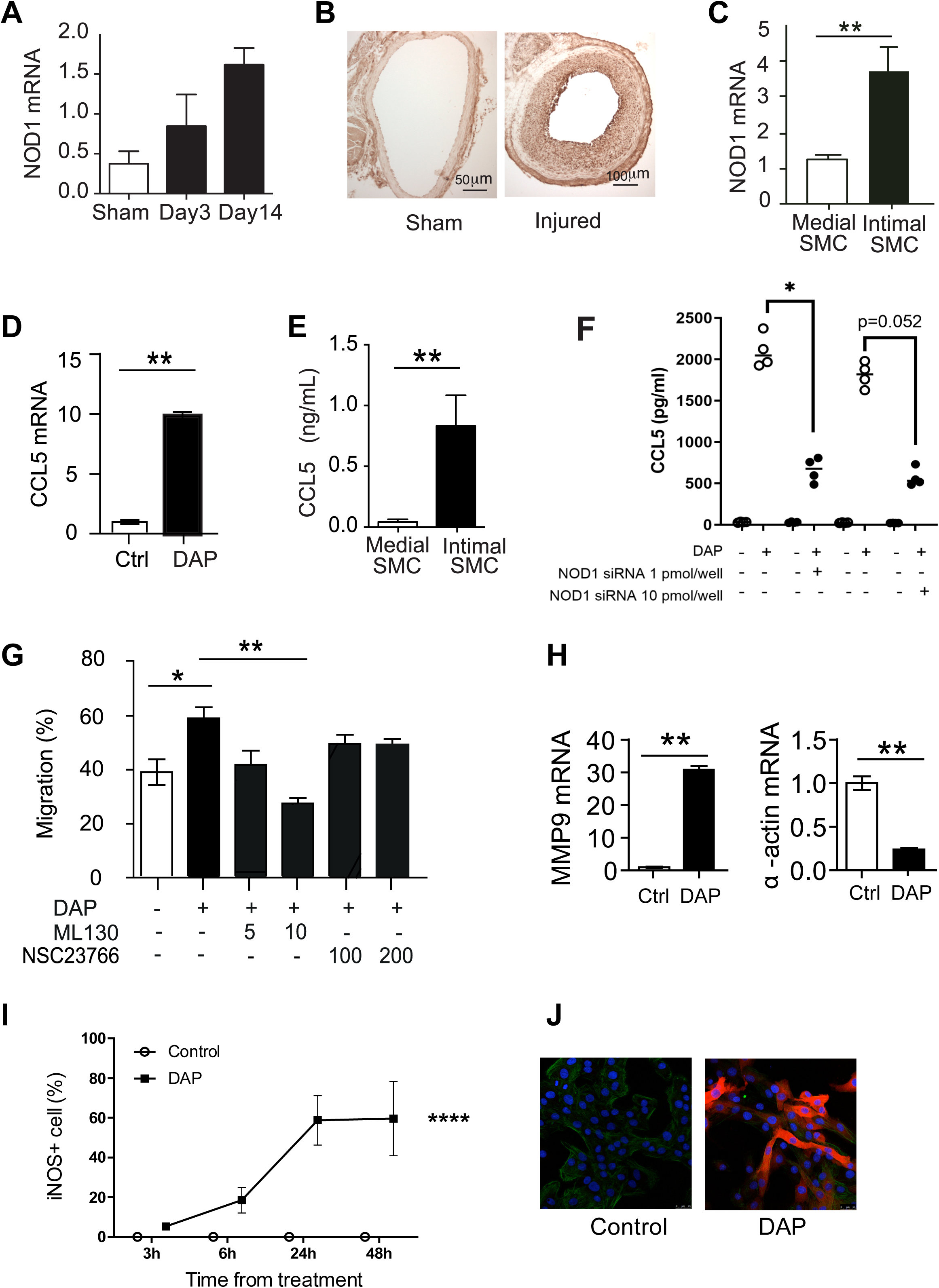
Intimal SMC function as NOD1 effector cells. (A) qPCR analysis of NOD1 expression in rat carotid arteries 3 days and 14 days after balloon injury. (B) Representative micrographs showing immunostaining of NOD1 in rat carotid arteries 14 days after balloon injury. (C) qPCR analysis of NOD1 levels in rat primary SMC isolated from media of uninjured carotid artery (Medial SMC) or from neointima 14 days after balloon injury (Intimal SMC). (D) qPCR analysis of *CCL5* in rat intimal SMC treated with or without C12-iE-DAP (DAP, 1µg/ml) for 24 h. (E) ELISA assessment of CCL5 production by the SMCs as in (C) in the absence exogenous stimulation for 24 h. (F) silencing RNA (siRNA) towards NOD1 receptor (G) Wound healing assay of rat intimal SMC migration in response to C12-iE-DAP (DAP, 1 µg/ml) and in the presence or absence of NOD1 inhibitor ML130 (5 µM or 10 µM) or small Rho GTPase Rac1 inhibitor NSC23766 (100 µM or 200 µM). (H) qPCR analysis of *MMP9* and smooth muscle α*-actin* mRNA (SMA) expression in rat intimal SMC treated with or without C12-iE-DAP (DAP, 1µg/ml) for 24 h.(I) time curve of iNOS expressing intimal SMC treated with or without DAP (J) representative micrographs showing iNOS immunofluorescence staining in cells from (I) at 24 h. Data are presented as mean ± SEM. Mann-Whitney test was performed, *p< 0.05, **p < 0.01.

#### Medal vs intimal SMC

*In vitro* studies of neointima derived NOD1^high^ SMC revealed that these SMC responded strongly to tri-DAP, the cognate NOD1 agonist discovered in peptidoglycan^4^. Tri-DAP stimulation increased CCL5 mRNA levels (Figure 7D), one of the chemokines induced *in vivo* in response to FK565. Another feature of NOD1^high^ SMC was their spontaneous release of CCL5 in the absence of exogenous stimulation (Figure 7E). Furthermore, when using a siRNA approach to knock-down the NOD1 expression, this significantly decreased the DAP mediated CCL5 increase (Figure 7F).

Interestingly, in an *in vitro* wound healing assay, NOD1 ligand tri-DAP promoted NOD1^high^ SMC migration, whereas NOD1 inhibitor ML130 decreased migration (Figure 7G). However, migration was not dependent on small ras homologous (Rho) nucleotide guanosine triphosphate-(GTP)ase ras-related C3 botulinum toxin substrate (Rac)1 given that blockade with Rac1 inhibitors did not influence migration. Moreover, NOD1 ligand induced expression of matrix metalloproteinase 9 and attenuated expression of smooth muscle α-actin in NOD1^high^ SMC (Figure 7H), as well as, a time-dependent increase in iNOS levels with NOD1 agonist (Fig. 7I-J).

## 4. Discussion

NOD1 ligation can enhance vascular inflammation and atherosclerosis in experimental animals^10,11^ but the cellular targets and responses mediating these responses have remained unclear. Here we report that a population of NOD1^high^ SMC is present in human and murine atherosclerotic lesions. NOD1 ligation of such cells triggered a set of proinflammatory responses involving MAPK and NF-kB activation, adhesion molecule expression, and production of proinflammatory cytokines and chemokines. *In vivo*, NOD1 ligands instigated transvascular inflammation, provoked intimal hyperplasia, accelerated atherogenesis and promoted coronary artery obstruction in experimental animals. The proatherogenic effect of NOD1 ligation was unaffected by NOD1 ablation of myeloid cells, implying that it requires vascular NOD1 activation. Together, these findings identify an important, proinflammatory and proatherosclerotic role of SMC responding to innate immune signals.

A few studies have investigated the expression and function of NOD1 in human atherosclerosis. Recently, NOD1 was described to be expressed in the intima of human coronary atherosclerosis ^15,16^ yet, the functional role in human atherosclerosis remains undefined. We confirm the expression of NOD1 in human coronary and carotid plaques as well as showing an increased expression of NOD1 in patients with symptomatic carotid plaques. Interestingly, NOD1 expression is mainly co-localized with α-actin positive SMCs in both the atherosclerotic lesions and the media layer of the vessel wall. Furthermore, using an *ex vivo* tissue culture system^8,17^ we demonstrated that NOD1 stimulation activates MAP kinases as well as ERK signaling, resulting in increased cytokine production. Thus, NOD1 activation could contribute to the increased inflammatory milieu in human atherosclerotic plaques, and possibly also contribute to a less stable plaque phenotype.

The present study demonstrates that NOD1 agonist, orally administered at high concentration, has potent effects on both innate and adaptive immunity. This is consistent with previous studies demonstrating the proatherogenic effect of NOD1 stimulation^12^. In addition to signs of macrophage and Th1-cell activation, NOD1 stimulation enhanced vascular IL-17 mRNA levels. IL-17 could contribute to several key mechanisms in NOD1 mediated vascular inflammation and atherogenesis, such as neutrophil infiltration^18^, SMC migration and collagen production^19,20^.

Another prominent property of NOD1 stimulation was the induction of chemokines. FK565-treatment led to 40-fold increase of CLL2, CCL5 and CXCL10 in arteries of mice. This coincided with signs of increased leukocyte infiltration, production of inflammatory cytokines and MMPs as well as impaired vessel wall integrity. The increased chemokine production is a feature of NOD1 stimulation also seen by others^11,12^. These observations support a concept where NOD1 ligand elicits an inflammatory cascade, initiated by local upregulation of chemokines and subsequent transvascular recruitment and activation of Ly6G^+^ neutrophils, CD68^+^ macrophages and CD3^+^ T lymphocytes, contributing to vascular injury. The NOD1 induced increase in MMP expression has been reported previously ^11^ and we can here demonstrate that this MMP activation also coincides with a decreased α-actin expression and elastic lamellae destruction, all hallmarks of vascular wall remodeling^21^ and plaque vulnerability ^22^.

From the bone-marrow transplantation study we can conclude that myeloid NOD1 does not promote atherosclerosis under unstimulated conditions, thus, the major effector cell for NOD1 stimulation is not of hematopoietic origin, thereby confirming previous reports^12,16^. However, the findings in our NOD1 treated mice mirrors the findings in ApoE−/− mice lacking NOD1, ie. ApoE−/− / NOD1−/− mice display decreased atherosclerosis, decreased immune cell infiltration and increased α-SMC^+^ cells in the lesions^15,16^. Although, there was no difference in proliferation in the atherosclerotic lesions, the decreased apoptotic activity in ApoE−/− mice lacking NOD1 further supports a role for NOD1 in regulating plaques stability^15,16^.

Previous studies have reported an important role of NOD1 expression in endothelial cells, mediating an up-regulation of adhesion molecule VCAM-1^16^. Interestingly, the current study instead highlights the importance of SMCs, and more specifically, a sub-population of SMC with imprinted high NOD1expression both in the artery of rodents and humans. Most NOD1^high^ SMC express vascular smooth muscle cell conserved filament protein alpha smooth muscle actin (α-SMA) but are distinguished from conventional medial SMC by constitutive twofold higher mRNA level of NOD1.

The NOD1^high^ SMC were highly responsive to exogenous NOD1 agonist and activated NF-κB, was associated with NOD1^high^ SMC in atherosclerotic lesion of *Ldlr*^−/−^ mice treated with NOD1 ligand. In addition to bacterial peptidoglycan derived NOD1 ligand, NOD1^high^ SMC also responds to endogenous danger associate molecular patterns (DAMP) derived from vascular injury. Indeed, our previous study in the rat model of vascular injury showed that activated NF-κB signal was present in approximately 80% of intimal SMC. Moreover, targeting IκB kinase β (IKKβ) in the injured artery could attenuate vascular inflammatory responses and neointima development^23^. The present study shows that a majority of cells in the injury-induced intimal lesion are NOD1^high^ SMC. Given these similarities, the NF-κB positive intimal SMC and injury responsive NOD1^high^ SMC are very likely overlapping populations. Emerging evidence shows that NOD1 is involved in transduction of danger signal of endoplasmic reticulum (ER) stress via activation of inositol-requiring enzyme 1α-TNF receptor-associated factor 2 (IRE1α/TRAF2) pathway^24^, implicating NOD1^high^ SMC function in several inflammatory situations. Besides ER stress, HMGB1 released, as a consequence of cell death or actively secreted by living cells under stress, is another known danger signal that correlates closely with severity of injuries. Furthermore, generation of other danger signals such as heat shock proteins has been reported in the context of vascular injury. In addition, also insulin and oxLDL has been demonstrated to activate NOD1 in vitro ^15,25^. The exact danger signals involved in vascular injury that are recognized by NOD1^high^ SMC remain to be elucidated in future studies.

NOD1^high^ SMC have strong inflammatory activity, as demonstrated by the release of CCL5 in the absence of extra stimulation. In a study on *Apoe*^−/−^ mice, Kanno et al. showed that CCL5 protein was present in the intima layer of arteries^12^. Thus, both *in vitro* and *in vivo* observations point to NOD1^high^ SMC as a producer of CCL5^12^. In a previous study, we also found that intima-derived SMC are characterized by high expression of a spectrum of chemokines including those highly expressed in the vessels of NOD1 ligand treated mice from the current study^26^. These findings in combination suggest that NOD1^high^ SMC have considerable capability in producing chemokines, thereby contributing to NOD1-induced transvascular inflammation and accelerated atherosclerosis.

Interestingly, NOD1^high^ SMC was the predominant cellular component of the intimal lesions found in both the stenotic atherosclerotic lesions in the mouse study, as well as, in the rat balloon injury model, a model if intimal hyperplasia. Given that NOD1 stimulation can induce a proliferative phenotype in vascular SMCs ^25^, this suggests that NOD1^high^ SMC drive intimal thickening, arterial narrowing and progressive occlusion in response to inflammatory or mechanical injury. Exploring the mechanisms behind these effects, we found that NOD1 agonist *per se* could promote SMC migration, since this effect was abolished when using NOD1 inhibitor, and subsequently promoting SMC MMP9 expression, downregulation of SMC α-actin and a time-dependent increase in iNOS^+^ cells. Thus, NOD1^high^ SMC are prominently contributing to vascular wall remodeling.

Notably, NOD1 agonist FK565 caused a 60% mortality. Oral administration of NOD1 agonist have previously been reported to cause septic shock and organ dysfunction^9^ and this cannot be ruled out in the current study. However, no apparent pathological changes of liver or kidneys were identified in the necropsies by gross examination.

An important question for future studies is the origin of NOD1^high^ SMC. Elegant lineage tracking studies in mouse arteries in scenarios of atherosclerosis, injury, and inflammation have established a consensus that majority of intimal SMC are derived from tunica media^27,28^. Consistent with this view, tracking NOD1-expressing SMC we found a small number of NOD^high^ SMC residing in the tunica media of arteries in *Ldlr^−/−^* mice.

In summary, these findings suggest that a subpopulation of vascular NOD1^high^ SMC in humans and rodents can initiate transvascular inflammation and lesion development in response to bacterial motifs and/or vascular injury. Understandings of where, when, and how the NOD1^high^ SMC are activated may open up possibilities for new therapeutic interventions.

## Acknowledgements

We thank Linda Haglund and Anneli Olsson for their technical assistance. We thank Astellas Pharma Inc for providing FK565.

## References

1. Gistera A, Hansson GK. The immunology of atherosclerosis. Nat Rev Nephrol. 2017;13:368–380. doi: 10.1038/nrneph.2017.51

2. Libby P, Hansson GK. From Focal Lipid Storage to Systemic Inflammation JACC Review Topic of the Week. Journal of the American College of Cardiology. 2019;74:1594–1607. doi: 10.1016/j.jacc.2019.07.061

3. Chen G, Shaw MH, Kim YG, Nunez G. NOD-like receptors: role in innate immunity and inflammatory disease. Annu Rev Pathol. 2009;4:365–398. doi: 10.1146/annurev.pathol.4.110807.092239

4. Chamaillard M, Hashimoto M, Horie Y, Masumoto J, Qiu S, Saab L, Ogura Y, Kawasaki A, Fukase K, Kusumoto S, et al. An essential role for NOD1 in host recognition of bacterial peptidoglycan containing diaminopimelic acid. Nat Immunol. 2003;4:702–707. doi: 10.1038/ni945

5. Inohara N, Ogura Y, Fontalba A, Gutierrez O, Pons F, Crespo J, Fukase K, Inamura S, Kusumoto S, Hashimoto M, et al. Host recognition of bacterial muramyl dipeptide mediated through NOD2. Implications for Crohn’s disease. J Biol Chem. 2003;278:5509–5512. doi: 10.1074/jbc.C200673200

6. Ogura Y, Inohara N, Benito A, Chen FF, Yamaoka S, Nunez G. Nod2, a Nod1/Apaf-1 family member that is restricted to monocytes and activates NF-kappaB. J Biol Chem. 2001;276:4812–4818. doi: 10.1074/jbc.M008072200

7. Inohara N, Koseki T, del Peso L, Hu Y, Yee C, Chen S, Carrio R, Merino J, Liu D, Ni J, Nunez G. Nod1, an Apaf-1-like activator of caspase-9 and nuclear factor-kappaB. J Biol Chem. 1999;274:14560–14567.

8. Johansson ME, Zhang XY, Edfeldt K, Lundberg AM, Levin MC, Boren J, Li W, Yuan XM, Folkersen L, Eriksson P, et al. Innate immune receptor NOD2 promotes vascular inflammation and formation of lipid-rich necrotic cores in hypercholesterolemic mice. Eur J Immunol. 2014;44:3081–3092. doi: 10.1002/eji.201444755

9. Cartwright N, Murch O, McMaster SK, Paul-Clark MJ, van Heel DA, Ryffel B, Quesniaux VF, Evans TW, Thiemermann C, Mitchell JA. Selective NOD1 agonists cause shock and organ injury/dysfunction in vivo. American journal of respiratory and critical care medicine. 2007;175:595–603. doi: 10.1164/rccm.200608-1103OC

10. Moreno L, McMaster SK, Gatheral T, Bailey LK, Harrington LS, Cartwright N, Armstrong PC, Warner TD, Paul-Clark M, Mitchell JA. Nucleotide oligomerization domain 1 is a dominant pathway for NOS2 induction in vascular smooth muscle cells: comparison with Toll-like receptor 4 responses in macrophages. Br J Pharmacol. 2010;160:1997–2007. doi: 10.1111/j.1476-5381.2010.00814.x

11. Nishio H, Kanno S, Onoyama S, Ikeda K, Tanaka T, Kusuhara K, Fujimoto Y, Fukase K, Sueishi K, Hara T. Nod1 ligands induce site-specific vascular inflammation. Arterioscler Thromb Vasc Biol. 2011;31:1093–1099. doi: 10.1161/ATVBAHA.110.216325

12. Kanno S, Nishio H, Tanaka T, Motomura Y, Murata K, Ihara K, Onimaru M, Yamasaki S, Kono H, Sueishi K, Hara T. Activation of an innate immune receptor, Nod1, accelerates atherogenesis in Apoe−/− mice. J Immunol. 2015;194:773–780. doi: 10.4049/jimmunol.1302841

13. Folkersen L, Persson J, Ekstrand J, Agardh HE, Hansson GK, Gabrielsen A, Hedin U, Paulsson-Berne G. Prediction of ischemic events on the basis of transcriptomic and genomic profiling in patients undergoing carotid endarterectomy. Mol Med. 2012;18:669–675. doi: 10.2119/molmed.2011.00479

14. Karadimou G, Folkersen L, Berg M, Perisic L, Discacciati A, Roy J, Hansson GK, Persson J, Paulsson-Berne G. Low TLR7 gene expression in atherosclerotic plaques is associated with major adverse cardio- and cerebrovascular events. Cardiovasc Res. 2017;113:30–39. doi: 10.1093/cvr/cvw231

15. Gonzalez-Ramos S, Fernandez-Garcia V, Recalde M, Rodriguez C, Martinez-Gonzalez J, Andres V, Martin-Sanz P, Bosca L. Deletion or Inhibition of NOD1 Favors Plaque Stability and Attenuates Atherothrombosis in Advanced Atherogenesis (dagger). Cells. 2020;9. doi: 10.3390/cells9092067

16. Gonzalez-Ramos S, Paz-Garcia M, Rius C, Del Monte-Monge A, Rodriguez C, Fernandez-Garcia V, Andres V, Martinez-Gonzalez J, Lasuncion MA, Martin-Sanz P, et al. Endothelial NOD1 directs myeloid cell recruitment in atherosclerosis through VCAM-1. FASEB J. 2019;33:3912–3921. doi: 10.1096/fj.201801231RR

17. Liu HQ, Zhang XY, Edfeldt K, Nijhuis MO, Idborg H, Back M, Roy J, Hedin U, Jakobsson PJ, Laman JD, et al. NOD2-mediated innate immune signaling regulates the eicosanoids in atherosclerosis. Arterioscler Thromb Vasc Biol. 2013;33:2193–2201. doi: 10.1161/ATVBAHA.113.301715

18. Geddes K, Rubino SJ, Magalhaes JG, Streutker C, Le Bourhis L, Cho JH, Robertson SJ, Kim CJ, Kaul R, Philpott DJ, Girardin SE. Identification of an innate T helper type 17 response to intestinal bacterial pathogens. Nat Med. 2011;17:837–844. doi: 10.1038/nm.2391

19. Taleb S, Tedgui A, Mallat Z. IL-17 and Th17 cells in atherosclerosis: subtle and contextual roles. Arterioscler Thromb Vasc Biol. 2015;35:258–264. doi: 10.1161/ATVBAHA.114.303567

20. Gistera A, Robertson AK, Andersson J, Ketelhuth DF, Ovchinnikova O, Nilsson SK, Lundberg AM, Li MO, Flavell RA, Hansson GK. Transforming growth factor-beta signaling in T cells promotes stabilization of atherosclerotic plaques through an interleukin-17-dependent pathway. Sci Transl Med. 2013;5:196ra100. doi: 10.1126/scitranslmed.3006133

21. Renna NF, de Las Heras N, Miatello RM. Pathophysiology of vascular remodeling in hypertension. Int J Hypertens. 2013;2013:808353. doi: 10.1155/2013/808353

22. Bentzon JF, Otsuka F, Virmani R, Falk E. Mechanisms of plaque formation and rupture. Circ Res. 2014;114:1852–1866. doi: 10.1161/CIRCRESAHA.114.302721

23. Bu DX, Erl W, de Martin R, Hansson GK, Yan ZQ. IKKbeta-dependent NF-kappaB pathway controls vascular inflammation and intimal hyperplasia. FASEB J. 2005;19:1293–1295. doi: 10.1096/fj.04-2645fje

24. Keestra-Gounder AM, Byndloss MX, Seyffert N, Young BM, Chavez-Arroyo A, Tsai AY, Cevallos SA, Winter MG, Pham OH, Tiffany CR, et al. NOD1 and NOD2 signalling links ER stress with inflammation. Nature. 2016;532:394–397. doi: 10.1038/nature17631

25. Shiny A, Regin B, Mohan V, Balasubramanyam M. Coordinated augmentation of NFAT and NOD signaling mediates proliferative VSMC phenotype switch under hyperinsulinemia. Atherosclerosis. 2016;246:257–266. doi: 10.1016/j.atherosclerosis.2016.01.006

26. Zhang C, Yang J, Jennings LK. Attenuation of neointima formation through the inhibition of DNA repair enzyme PARP-1 in balloon-injured rat carotid artery. Am J Physiol Heart Circ Physiol. 2004;287:H659–666. doi: 10.1152/ajpheart.00162.2004

27. Shankman LS, Gomez D, Cherepanova OA, Salmon M, Alencar GF, Haskins RM, Swiatlowska P, Newman AA, Greene ES, Straub AC, et al. KLF4-dependent phenotypic modulation of smooth muscle cells has a key role in atherosclerotic plaque pathogenesis. Nat Med. 2015;21:628–637. doi: 10.1038/nm.3866

28. Feil S, Fehrenbacher B, Lukowski R, Essmann F, Schulze-Osthoff K, Schaller M, Feil R. Transdifferentiation of vascular smooth muscle cells to macrophage-like cells during atherogenesis. Circ Res. 2014;115:662–667. doi: 10.1161/CIRCRESAHA.115.304634

