## Supplementary material for "NOD1 ligand FK565 promotes atherogenesis and accumulation of NOD1^high^ smooth muscle cells in atherosclerotic lesions": Suppl Methods

**Materials and methods**

***Mouse treatment***

Female low-density lipoprotein receptor deficient (*Ldlr^-/-^*) mice were purchased from Jackson Laboratory (Bar Harbor, Maine, USA) and raised in a pathogen-free barrier facility. Water and food were available *ad libitum*. At the age of 11 weeks, *Ldlr^-/-^* mice were fed high-fat diet (1.25% cholesterol, D12108, Research Diets) and administrated with 21 µg/ml (estimated dose of 120µg/day/mouse, a kind gift from Astellas, Tsukuba, Japan) in drinking water *ad libitum* for 10 weeks (n=10 per group). Littermates receiving regular water was used as control. Mice were euthanized with carbon dioxide and blood was collected by cardiac puncture. The heart and vascular tree were then perfused with PBS. The aortic root was preserved for immunohistochemistry. All procedures involving mice were in accordance with the ARRIVE guidelines and approved by the Regional Northern Stockholm Animal Ethics Committee, in accordance with the European Communities Council Directives of 22 September 2010 (2010/63/EU).

***Bone marrow transplantation***

Male *Ldlr^-/-^* and *Nod1*^-/-^ mice and their corresponding C57Bl/6J controls were purchased from Jackson Laboratory (Bar Harbor, Maine, USA). Bone marrow transplantation was performed as previously described [1]. In brief, 6–9 week-old *Ldlr^-/-^* mice were irradiated with 2 doses of 700 rad 3 h apart and randomly assigned to receive bone marrow from *Nod1*^-/-^ or corresponding age-matched C57BL/6J control donor mice (n=10–11 recipients per group). The transplanted recipients were allowed to recover from the irradiation for 4 weeks, after which they were fed a high-fat diet (HCD, D12108, Research Diets, New Brunswick, NJ) for 8 weeks and then sacrificed.

***Lesion quantification***

As described previously [2], aortic arch was collected by micro-dissection to remove connective tissue and adventitia, fixed by 4% formaldehyde, and pinned out and stained with Sudan IV solution for 5 minutes. Aortic root was used for cryosection. 10 μm sections were collected starting at a 100-μm distance from the appearance of the aortic valves until 800 μm distance. Sections were air-dried and fixed with 4% formaldehyde in PBS or acetone for 10 min. Formaldehyde-fixed sections were stained with hematoxylin and oil red-O at 100-μm intervals. Innominate artery was applied for paraffin section. 5 μm sections were collected starting at branches of right common carotid artery and right subclavian artery until the aortic arch. Sections with biggest lesions were selected for immunostaining.

Images were captured for each section with a Leica DM-LB2 microscope equipped with a Leica DC300 camera and quantified using Leica Q-win or Image J.

***Histological analysis***

In mouse experiment, primary antibodies to CD68 (1:20 000, AbD Serotec, Düsseldorf, Germany), CD4 (1:100), Ly6G (1:800, clone 1A8, BD Biosciences, San Diego, CA), CD3 (1:300, AbD Serotec) and α-actin (1:1000, Abcam, Cambridge, UK) were applied to acetone-fixed aortic root cryosections; NOD1 (1: 150, Acris antibodies, Hiddenhausen, Germany), NF-κB (1:50, Cell Signaling) and α-actin (1:1000, Abcam, Cambridge, UK) were applied to innominate artery paraffin sections, followed by detection with the ABC alkaline phosphatase kit (Vector Laboratories, Burlingame, CA, USA) or Alex 488/594-conjugated secondary antibody. Elastin was examined by Verhoeff-Van Gieson (VVG) staining on formaldehyde-fixed aortic root sections and expressed as total number of destructed valves for each treatment. Elastin were stained black, and nuclear were blue-black.

In rat experiment, paraffin-embedded rat carotid artery sections were de-paraffinized and then rehydrated through grade ethanol washes. After incubation with rabbit anti-NOD1 antibody (Acris antibodies) at 4°C overnight, sections were incubated with biotinylated goat anti-rabbit followed by avidin-biotin peroxidase complex and developed with diaminobenzidine (all from Vector Laboratories, Burlingame, California, USA).

In human coronary artery (n=5) and human carotid plaque staining (n=14), NOD1 (1:100, R&D Systems) and α-actin (1:1000, Abcam, Cambridge, UK) were applied to paraffin sections, followed by detection with Alex 488/594-conjugated secondary antibody. Normal rabbit IgG was used as control for the NOD1 antibody. Co-localization of NOD1^high^SMC in human carotid plaques were detected by double immunofluorescence staining and assessed by manual quantification. Images were acquired using confocal microscopy. For each plaque, 1–5 regions of interest were analyzed, and the average per plaque was calculated. Data are expressed as percentage of NOD1^high^SMC of all α-actin^+^SMC and as percentage of NOD1^high^SMC of all NOD1 positive cells. Quantification and co-localisation of NOD1^high^SMC in the innominate artery of NOD1 ligand treated and saline treated mice was performed using ImageJ with the automated plugin JACoP. A threshold value was set manually for each channel, defined as the minimum pixel intensity considered to be signal and not background or noise. Correlation is presented as Manders Coefficients; M1 and M2, where M1 represents the ratio of red pixels that coincides with green pixels and vice versa for M2 All quantification was performed in a blinded manner.

***RNA isolation, cDNA synthesis and real-time PCR***

RNA was isolated using the RNeasy Kit (Qiagen Inc., Hilden, Germany). RNA quality was determined by capillary electrophoresis on a BioAnalyzer (Agilent Technologies, Waldbronn, Germany). cDNA was synthesized using random hexanucleotide primers and Superscript III reverse-transcriptase (Invitrogen Life Technologies, Paisley, UK).

Real-time PCR was performed on an ABI 7900HT Sequence Detector (Applied Biosystems, Foster City, CA). Mouse CCL5, CXCL10, CD68, CD3, CD4, CD8, IL6, IL17A, house-keeping gene hypoxanthine-guanine phosphoribosyltransferase (HPRT) were detected using Assays-on-Demand primers and probes (ABI, Foster City, CA). Mouse MMP9, MMP10, MMP12, IFNγ, MCP-1, IL10, IL12p35, IL12p40, IL1β, TNFα, iNOS, Arg-1, HO-1, house-keeping gene glyceraldehyde-3-phosphate dehydrogenase (GAPDH) and rat NOD1, CCL5, MMP9, ACTA2, house-keeping gene β-actin were analyzed using SYBRGreen primers (Invitrogen Life Technologies, Paisley, UK). We verified that the SYBRGreen primers amplified a single product with a distinct melting curve and a single specific band with the correct size by electrophoresis on agarose gels.

Relative mRNA was expressed in arbitrary units, calculated as 2^-ΔΔCT^, where ΔCT was the CT of the target gene after subtracting the CT value of the reference gene and ΔΔCT was the CT value corrected by the average CT of each group. The primer sequences were presented in Table S2.

***Blood and lipid analysis***

Whole blood anti-coagulated with EDTA was analyzed on a Scil Vet abc hemocounter (Scil animal care company, Gurnee, IL). Cholesterol and triglycerides in the plasma were measured using enzymatic colorimetric kits (Randox Laboratories, London, UK) following manufacture’s instruction. Data were presented as median (25% percentile-75% percentile), Mann-Whitney test.

***Rat intima smooth muscle cell culture***

Male Sprague-Dawlery rats were subjected to angioplasty injury to left common carotid artery to de-endothelial under general anesthetization by an intraperitoneal injection of pentobarbital (2mg/kg) plus Hypnorm^®^(50mg/kg, Janssen Pharmaceutica, Belgium) as previously described [4]. Briefly, a Fogarty F2 balloon catheter (Baxter Healthcare Corp., Round Lake, IL, USA) was introduced through the left external carotid artery and advanced into the common carotid artery; the balloon was inflated with 0.15 ml saline and then withdrawn to the entry point. The entire procedure was repeated three times. Animal were killed by overdosing pentobarbital at day 3 and day 14 after injury and perfused with 100 ml phosphate-buffered saline (PBS) via the left ventricle before carotid arteries were harvested.

SMCs were isolated from the medial layer of the thoracic aorta of 6-week-old male Sprague-Dawley rats, and intimal SMCs were derived from the carotid artery 2 weeks after balloon angioplasty. Cells were maintained under standard cell culture conditions (+37°C, 5% CO2). Rat medial and intimal SMCs were grown in Dulbecco’s Modified Eagles medium (DMEM, invitrogen Corporation) supplement with 10% (vol/vol) heat-inactivated fetal bovine serum (FCS), 1 mmoll/l L-glutamine and antibiotics (penicillin G 100 U/ml, and streptomycin 100 μg/ml, Gibco BRL).

***siRNA experiments***

The scramble siRNA (4390843) and NOD1 siRNA (s178978) were purchased from life technology for the RNA interference study in rat medial SMCs. Following the manufacturer's instructions, the cellular delivery of 1 pmol and 10 pmol siRNA (per well) was carried out using Lipofectamine MAXi (Invitrogen 13778-030, CA, USA). Transfections were allowed to proceed for 24 hours followed by DAP (1µg/ml) stimulation for 24 hours, after which the supernatants and cells were collected and kept at -80 degrees.

***iNOS expression in rat intimal SMCs and quantification***

INOS expression in rat intimal SMCs was detected by immunofluorescence staining using an anti-iNOS antibody (N32030, BD Biosciences, 1:50). Images were acquired using confocal microscopy. INOS-positive cells were manually counted in five randomly selected areas per slide, and the percentage of positive cells was calculated.

***Wound healing assay***

In wound healing assay, rat intima SMCs culture conditions were optimized to ensure homogeneous and viable cell monolayers prior to wounding. When the cell confluence reached ~90%, cells were treated with medium containing 5mmol/L hydroxyurea to inhibit cell proliferation for 24h before a homogenous artificial wound was created using a sterile plastic 10 μl micropipette tip and cell debris was removed by washing the cells in serum-free medium. Cells were then treated with C12-iE-DAP (1µg/ml) for 24h with or without GTPase Rac1 inhibitor NSC23766 or NOD1 inhibitor ML130. The area between the borders of the wound was photographed using inverted phase contrast microscope and quantified before and after the cell migration.

***Expression phenotype correlation***

NOD1 expressions in symptomatic patients were compared to asymptomatic patients as previously described [5]. In brief, patients diagnosed with >70% carotid stenosis were referred to Karolinska Hospital, Stockholm, Sweden for surgical treatment. The investigation was approved by the Ethical Committee of Northern Stockholm. Carotid plaques from endarterectomy were immediately rinsed in ice-cold sterile saline and instantly frozen on dry ice for later isolation of RNA using RNeasy total RNA isolation kit (Qiagen). Affymetrix HG-U133A Genechip arrays were performed on 10µg total RNA. Gene expressions were compared between symptomatic versus asymptomatic patients using in-house software (KIGeneConnect).

***Human carotid plaque tissue culture***

Seven carotid artery plaques were collected from patients undergoing endarterectomy at Karolinska University Hospital, Stockholm, Sweden. The investigation was approved by the Ethical Committee of Northern Stockholm. Human carotid plaque tissue cultures were set up as previously described [6]. In brief, fresh plaque tissues were processed to remove calcified tissue, cut into small pieces (about 1.5 mm^3^), and washed with cold PBS. The tissue was distributed equally in a 48-well plate (0.1 g tissue/well). One hour after incubation in RPMI with 10% FCS, MAPK p38 inhibitor SB 203580, MEK 1 inhibitor PD 98059, JNK Inhibitor II SP600125 (Merck Chemicals Ltd, Nottingham, UK), NF-kB inhibitor BAY-11-7082 (Cayman, Ann Arbor, MI, USA), and lauroyl-γ-D-glutamyl-meso-diaminopimelic acid (C12-iE-DAP, Invivogen, Sweden) were administered for the indicated time. Supernatants and plaque tissue were snap-frozen and kept at -80 ^o^C until further analysis.

***Western blot***

Human carotid plaque tissue was lysed in tissue lysis buffer (Thermo Fischer Scientific, Rockford, IL) containing protease and phosphatase inhibitor cocktail (Thermo Fischer Scientific). Protein concentration was measured by BCA assay (Bio-Rad, Hercules, CA). Equal amounts of proteins (50–80 mg) were mixed 1:1 (v/v) with sample buffer (Bio-Rad), and separated on a 12% SDS-PAGE gel, transferred to a polyvinylidene difluoride membrane, and probed with following antibodies: mouse anti-phospho-p38, rabbit anti-p38, rabbit anti-phospho-ERK, rabbit anti-EKR1/2, mouse anti-phospho-SAPK/JNK, rabbit anti-JNK, rabbit anti-phospho-IκBa (Cell Signaling, Danvers, MA), and rabbit anti-IκBa (clone C-15, Santa Cruz Biotechnology, CA). Phosphorylated and total proteins were probed on the same membrane using the Re-blot kit (Millipore, Billerica, MA) and visualized using ECL (Bio-Rad).

***Cytokine assays***

Human IL-1β, IL-6, IL-8, and IL-10 and rat CCL5 were measured using ELISA kits (R&D Systems, Minneapolis, MN and Mabtech, Nacka Strand, Sweden) according to the manufacturers’ instructions.

**Statistics**

Data are presented as mean ± SEM unless otherwise stated. Differences between groups were tested by Mann-Whitney U test, Chi-square test, Two-way ANOVA followed by Bonfferoni’s test, Wilcoxon matched-pairs test, with *P* values as reported. *P* value of <0.05 was considered significant.

**Table S1**. Non-fasting plasma triglycerides (TG), cholesterol (CHOL) levels and body weight (BW).

|  | TG (mmol/l) | CHOL (mmol/l) | BW (g) |
| --- | --- | --- | --- |
| **FK565 (21µg/ml) experiment** |  |  |  |
| Control | 3.5 (3.3-3.8) | 14.2 (12.4-15.7) | 20.7 ± 0.5 |
| FK565 (21µg/ml) | 3.0 (2.0-6.9) | 14.1 (13.6-17.0) | 19.0±0.6 |
| *P value* | *0.57* | *0.55* | *0.047* |
| **Bone marrow transplantation** |  |  |  |
| Wild-type chimeras | 4.0 (3.7-4.5) | 23.5 (21.6-25.8) | 23.7 ± 0.4 |
| Nod1-/- chimeras | 4.7 (4.0-6.1) | 24.6 (23.2-27.8) | 23.7±0.5 |
| *P value* | *0.15* | *0.32* | *0.97* |

**Table S2.** (A) Mouse primer sequences for real-time PCR

| *Gene* | Forward primer | Reverse primer |
| --- | --- | --- |
| *Il6* | TCCAGTTGCCTTCTTGGGAC | GTGTAATTAAGCCTCCGACTTG |
| *Tnf* | GGCTGCCCCGACTACGT | GACTTTCTCCTGGTATGAGATAGCAAA |
| *Ifng* | TGGGTTGACCTCAAACTTGGC | GGCCATCAGCAACAACATAAGCGT |
| *Ccl2* | TTAAAAACCTGGATCGGAACCAA | GCATTAGCTTCAGATTTACGGGT |
| *Il10* | GGTTGCCAAGCCTTATCGGA | ACCTGCTCCACTGCCTTGCT |
| *Gapdh* | AGGCCGGTGCTGAGTATGTC | TGCCTGCTTCACCACCTTCT |
| *Mmp9* | CTGTCCAGACCAAGGGTACAGCCT | GAGGTATAGTGGGACACATAGTGG |
| *Il12a* | ACCTGCTGAAGACCACAGATGACA | TAGCCAGGCAACTCTCGTTCTTGT |
| *Il12b* | CCAATTACTCCGGACGGTTC | AGTCCCTTTGGTCCAGTGTG |
| *Il1b* | TGGTGTGTGACGTTCCCATT | CAGCACGAGGCTTTTTTGTTG |
| *Nos2* | CAGCTGGGCTGTACAAACCTT | CATTGGAAGTGAAGCGTTTCG |
| *Arg1* | GTATGACGTGAGAGACCACG | CTCGCAAGCCAATGTACACG |
| *Hmox1* | CCTTCAAGGCCTCAGACAAA | GAGCCTGAATCGAGCAGAAC |

**Table S3.** Rat primer sequences for real-time PCR

| *Gene* | Forward primer | Reverse primer |
| --- | --- | --- |
| *Nod1* | GCTCATCCGGACCAAAACTA | CAATGTGTCGCTCTCCTTGA |
| *Nod2* | TCCCTGTCTTCTCATGGATGGTGT | AAGAGGTACATGTCCGTGCTGGTT |
| *Actb* | AGA GCG AAA TCG TGC GTG AC | CAA TAG TGA TGA CCT GGC CGT |
| *Ccl5* | GTGCCCACGTGAAGGAGTAT | TCTTCTCTGGGTTGGCACAC |
| *Cx3cl1* | CGGCATGACGAAATGCAACA | ATGGCGTCTTGGACCCATTT |

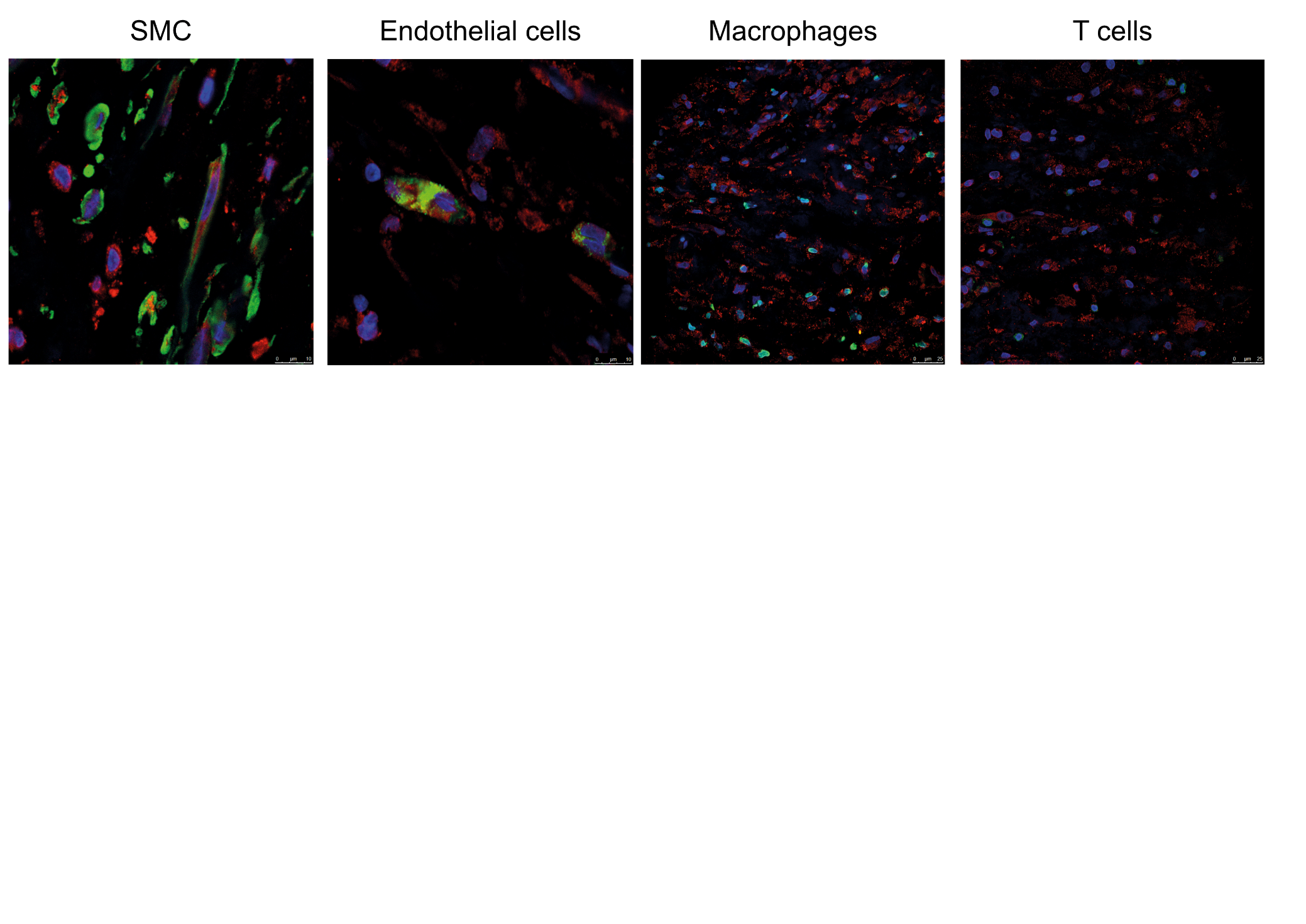

**Figure S1. Representative staining of human carotid atherosclerotic plaques.** *Representative IF staining of NOD1 (green) with markers for SMC, Endothelial cells, macrophages and T cells in human carotid plaques.*

**References**

1.	Lundberg, A.M., et al., *Toll-like receptor 3 and 4 signalling through the TRIF and TRAM adaptors in hematopoietic cells promotes atherosclerosis.* Cardiovasc Res, 2013.

2.	Nicoletti, A., et al., *Immunoglobulin treatment reduces atherosclerosis in apo E knockout mice.* J Clin Invest, 1998. **102**(5): p. 910-8.

4.	Bu, D.X., et al., *IKKbeta-dependent NF-kappaB pathway controls vascular inflammation and intimal hyperplasia.* FASEB J, 2005. **19**(10): p. 1293-5.

5.	Agardh, H.E., et al., *Expression of fatty acid-binding protein 4/aP2 is correlated with plaque instability in carotid atherosclerosis.* J Intern Med, 2011. **269**(2): p. 200-10.

6.	Liu, H.Q., et al., *NOD2-Mediated Innate Immune Signaling Regulates the Eicosanoids in Atherosclerosis.* Arterioscler Thromb Vasc Biol, 2013. **33**(9): p. 2193-2201.
